# Responses of *Escherichia coli* and *Listeria monocytogenes* to ozone treatment on non-host tomato: Efficacy of intervention and evidence of induced acclimation

**DOI:** 10.1101/2021.08.13.456288

**Authors:** Xiaomei Shu, Manavi Singh, Naga Bhushana Rao Karampudi, David F. Bridges, Ai Kitazumi, Vivian C. H. Wu, Benildo G. De los Reyes

## Abstract

Because of the continuous rise of foodborne illnesses caused by the consumption of raw fruits and vegetables, effective post-harvest anti-microbial strategies are needed. This study evaluated the dose × time effects on the anti-microbial action of ozone (O_3_) gas against the Gram-negative *Escherichia coli* O157:H7 and Gram-positive *Listeria monocytogenes,* which are common contaminants in fresh produce. The study on non-host tomato environment correlated the dose × time aspects of xenobiosis by examining the correlation between bacterial survival in terms of log-reduction and defense responses at the level of gene expression. In *E. coli*, low (1 μg O_3_/g of fruit) and moderate (2 μg O_3_/g of fruit) doses caused insignificant reduction in survival, while high dose (3 μg/g of fruit) caused significant reduction in survival in a time-dependent manner. In *L. monocytogenes*, moderate dose caused significant reduction even with short-duration exposure. Distinct responses to O_3_ xenobiosis between *E. coli* and *L. monocytogenes* are likely related to differences in membrane and cytoplasmic structure and components.

Transcriptome profiling by RNA-Seq showed that primary defenses in *E. coli* were attenuated after exposure to a low dose, while the responses at moderate dose were characterized by massive upregulation of pathogenesis and stress-related genes, which implied the activation of defense responses. More genes were downregulated during the first hour at high dose, with a large number of such genes getting significantly upregulated after 2 hr and 3 hr. This trend suggests that prolonged exposure led to potential adaptation. In contrast, massive downregulation of genes was observed in *L. monocytogenes* regardless of dose and exposure duration, implying a mechanism of defense distinct from that of *E. coli*. The nature of bacterial responses revealed by this study should guide the selection of xenobiotic agents for eliminating bacterial contamination on fresh produce without overlooking the potential risks of adaptation.

## Introduction

Perennial outbreaks of foodborne illnesses due to changes in pathogen population dynamics have an astonishing impact to human health [1–4]. In the United States alone, it was estimated that more than 30% of food-related deaths are due to the combined effects of only two bacterial pathogens, *i.e., Listeria monocytogenes* (28%) and *Escherichia coli* O157:H7 (3%) [5], both of which have been blamed for the more recent outbreaks on fresh vegetables including tomato (*Solanum lycopersicum* L.) [6, 7]. These pathogenic bacteria can survive under a wide range of environmental conditions and often contaminate their non-host plants at several developmental stages, and along the pre-harvest and post-harvest production pipelines through multiple routes [8, 9]. For instance, even a brief exposure of wounded tomato fruits to *E. coli* O157:H7 can provide an effective inoculum for widespread contamination during subsequent post-harvest handling and processing [10, 11].

Strategies including on-farm hygiene, decontamination by washing, film coating, prophage induction, and the use of chemical interventions that involve sodium hypochlorite (NaClO), sodium chlorite (NaClO_2_), acidified NaClO_2_, acidified sodium benzoate (NaB), or peracetic acid (PAA) are common means for post-harvest control of bacterial contamination on fresh tomatoes and other types of vegetables [12–14]. Ozone gas (O_3_) has been widely used decontaminating agent for eliminating *E. coli* O157:H7 and *L. monocytogenes* on various types of surfaces, including seeds, fresh fruits, as well as other organic and inorganic substrates such as milk and water [15–18]. However, the long-term impacts of such chemical intervention to potential acclimation, adaptation, and selection caused by chronic exposure to selective doses are often overlooked. For example, viable but non-culturable state of *E. coli* O157:H7 induced by exposure to various types of environmental stresses could contribute to adaptation [19]. The effects of environmental stresses including acid induced adaptation has also been shown to cause an acclimation effect in *L. monocytogenes* [20]. To address these concerns, comprehensive understanding of the potential consequences of sub-optimal, optimal, and supra-optimal doses and duration of exposure to chemical intervention is important in order to prevent future outbreaks caused by the proliferation of resistant strains triggered by acclimation to strong selective pressures [8].

Human-pathogenic bacteria respond to environmentally induced perturbations by employing a variety of defense or avoidance mechanisms [21, 22]. Bacterial defenses could be stimulated initially by different types of environmental factors, and such stimulation could further lead to a ‘priming effect’ that builds resistance to subsequent episodes of stress [23, 24]. For example, studies have shown that heat stress effectively primes *E. coli* O157 to develop resistance to subsequent exposure to acidic environments [25]. While surviving on non-host environments, such as the surfaces of fresh fruits and vegetables, bacterial cells are subjected to intense stress pressure, which could lead to acquired tolerance to other stresses or persistence as a viable inocula for much longer periods until they are revitalized in a suitable host environment [26]. Acquired resistance to environmental stresses could eventually promote anti-microbial resistance (AMR) through defense mechanisms that are effective against a broad range of chemical intervention agents. These AMR mechanisms also enable microorganisms to resist the anti-microbial effects of chemical agents, and such resistance has been suggested to be a major cause of perennial outbreaks in the food industry. For example, disinfectant-injured and genetically distinct sub-populations of *E. coli* believed to have originated from the widespread use of chlorine-based agents and O_3_ have been identified in water. There are also indications that these treatments trigger the proliferation of genetically distinct sub-populations that arise from the injurious but non-lethal effects of such chemical agents [27].

In our previous analysis of the transcriptional responses of *E. coli* O157:H7 to gaseous chlorine dioxide (ClO_2_) on non-host tomato environment, we uncovered characteristic gene expression signatures which indicated that optimal dose x time interaction causes an effective reduction of bacterial viability with evidence of injury and killing. However, gene expression signatures also indicated that supra-optimal dose x time effects could trigger resistance through acclimation or adaptation. Patterns in ClO_2_-mediated changes in gene expression revealed that longer exposure even under a dosage as low as 1 μg could effectively trigger new bursts of independent defense mechanisms on the surviving sub-populations of bacteria after the effective killing phase. These trends pointed to the occurrence of adaptation and selection in residual sub-populations surviving on the surface of non-host tomato [26]. Guided by these findings, we examined the effects of another commonly used chemical intervention agent, ozone (O_3_) gas on the Gram-negative *E. coli* O157:H7 and Gram-positive *L. monocytogenes* in context of the potential consequences of dose x time effects not only on the intervention efficacy but also on potential AMR on a common non-host environment, *i.e.,* fresh tomatoes. We discuss here the biological significance of the transcriptional networks associated with potential xenobiotic effects of O_3_ on *E. coli* and *L. monocytogenes*, and the implications of dose x time interaction to optimal and supra-optimal effects.

## Methods and Methods

### Microbial inocula

*E. coli* O157:H7 (ATCC 35150) and *L. monocytogenes* (ATCC 19115) were from the permanent cultures of the U.S. Department of Agriculture-Agricultural Research Service, Western Regional Research Center’s Pathogenic Microbiology Laboratory. Working cultures used throughout the study were maintained on tryptic soy agar (TSA) a 4°C and were inoculated from frozen stocks maintained at −80 °C in 25% glycerol broth. Prior to experimentation, the bacterial strains were cultured overnight at 37°C in tryptic soy broth (TSB), centrifuged at 5000xG for 15 min, re-suspended in 10 ml 0.1% peptone water, centrifuged for another 15 min and re-suspended in 12 ml 0.1% peptone water.

Samples of fresh tomato without post-harvest processing were obtained from Windset Farms, California. Tomato fruits without visual damage or mold growth were washed with water and, surface sterilized with 70% ethanol, and dried in the hood. Aliquots of 250 μl of bacterial suspension were inoculated on the surface of each tomato fruit, followed by air drying for a few hours. Inoculated fruits were placed in sterile plastic bags and incubated overnight at 4°C. Twelve tomato fruits were weighed (2.0 ± 0.1 kg) and used for each O_3_ treatment. Three replicates consisting of twelve tomato fruits in each replicate were included for each treatment. Tomato fruits were treated with 1 μg per g tomato fruits (low dose), 2 μg per g tomato fruits (moderate dose) or 3 μg per g tomato fruits (high dose) of O_3_ as previously described [8, 28].

### Regrowth assay on E. coli and L. monocytogenes

After 1 hr, 2 hr and 3 hr of O_3_ treatment, each tomato fruit that was inoculated with either *E. coli* or *L. monocytogenes* was rinsed with 10 ml 0.1% peptone water for 1 min for serial dilution (10^-1^ to 10^-5^) and plating using MacConkey Sorbital Agar supplemented with 0.05 mg/l Cefixime and 2.5 mg/L PotassiumTellurite (CT-SMAC) for *E. coli* or Polymyxin acriflavine lithium chloride ceftazidime aesculin mannitol (PALCAM) agar supplemented with 10 mg/l polymyxin B, 20 mg/l ceftazidime, and 5 mg/l acriflavine for *L. monocytogenes*. Both media were layered with a thin layer of TSA to aid in the recovery of sub-lethally injured cells. Samples were incubated overnight at 37°C and bacterial growth assay (log CFU/g) was determined by comparing O_3_-treated samples with the control according to standard procedures [8].

### RNA-Seq library construction, sequencing, and data processing

Bacterial RNA samples were isolated from each combination of dose (1 μg, 2 μg, 3 μg) and exposure duration (1 hr, 2 hr, 3 hr) in *E. coli* and *L. monocytogenes* using a Quick-RNA™ Fungal/Bacterial RNA Microprep kit (Zymo Research) according to the manufacturer’s instructions. For each sample, three replicates were used to construct three RNA-Seq libraries, which were then pooled and sequenced twice at 900x depth per library. Sequencing was performed on the Illumina HiSeq-3000 at 150-bp paired-end reads at the Genomics Core Facility, Oklahoma Medical Research Foundation, Norman, OK, USA.

Raw sequence reads from RNA-Seq were processed according to standard protocols [29]. Raw data were preprocessed with Cutadapt (v1.9.1) to remove adapters and low-quality sequences to generate paired 100-bp reads [30]. Subsequently, data with at least 16 million pairs per library were mapped using Edge-Pro (version v1.3.1) to account for polycistronic gene organization [31]. Reference *E. coli* O157:H7 str. Sakai genome (Genbank: GCA_000008865.2, NCBI: ASM886v2) and *L. monocytogenes* EGD-e (Genbank: GCA_000196035.1, NCBI: ASM19603v1) were used for mapping based on high map rates (∼98%) of control sample and availability of pathway annotation in KEGG [32].

### Propensity transformation and transcriptome analysis

Relative changes in gene expression expressed as Propensity Scores (PS) were established from the standard RPKM-based expression data using two batches of sequences for control (t_0_), 1 hr (t_1_), 2 hr (t_2_) and 3 hr (t_3_) for all three doses of O_3_. Average RPKM were transformed using the Propensity Transformation methodology that was optimized from the ClO_2_ study on *E. coli* [66] based on the following equation:

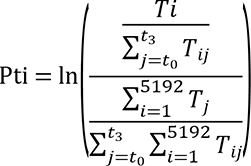

Where, *Pt_i_*=Propensity transformation of RPKM value of transcript *i*

*T_i_*= RPKM value of transcript *i*

*j*=Variable that iterates over datasets of *t_0_* =control, *t_1_*= 1 hr, *t_2_* = 2 hr and *t_3_* = 3 hr

*i*=Variable that iterates over the total number of transcript-encoding loci

Missing data in the dataset was considered as NULL and its transformed value was considered 0. The PS data from each library were assumed to form a normal distribution ranging between -n to +n, which were further fragmented into 20 quartiles based on the propensity scores. Quartile cuts resulted in 250 transcripts per quartile in all the datasets. Two quartiles per dataset representing the transcripts with the lowest and the highest propensity scores were subsequently selected for transcriptional regulatory network analysis, resulting 500 transcript-encoding loci per dataset (Additional file 3: Table S2). Two-way hierarchical clustering analysis with both RPKM values and PS values was performed using JMP, 11 (SAS Institute Inc., Cary, NC, USA).

### Genetic network modeling

A subset of normalized RPKM values were selected to calculate the standard Pearson Correlation Coefficient (PCC) using the Python Pandas library. The dataset was derived from the PS values without log transformation containing only positive values. The PCC for one versus all transcripts were calculated using this subset of normalized RPKM that resulted in a diagonally symmetrical matrix of 5129 × 5129 coefficients in which the diagonal values represent the PCC of every transcript locus with itself.

Transcript-encoding gene loci for network modeling were selected via two filtration steps: first using the PS followed by PCC. PS-based selection is as explained above using the quartile cuts that resulted in a group of transcript-encoding loci and their highly correlated co-expression partners were selected using the PCC values. The second step of transcript selection was based on PCC with cut-off of 0.9999 for filtration of positively and negatively correlated transcript loci. This selection represented the most likely O_3_-affected gene loci that were significant according to PS and their highly correlated co-upregulated, co-downregulated, or inversely co-expressed loci from the primary selection.

## Results

### Dose and time effects of O_3_ on Escherichia coli and Listeria monocytogenes

To understand the importance of dose × time dynamics on the xenobiotic action of O_3_ on foodborne bacterial pathogens, we compared the effects of chemical treatments on fresh tomato fruits between the Gram-negative *E. coli* O157:H7 and Gram-positive *L. monocytogenes* under a non-host environment provided by fresh tomatoes. We investigated the efficacy of bacterial killing by examining regrowth on fresh media using the washings from inoculated tomato fruits after different durations of treatment at different doses. Exposure to a low dose caused only mild effects on *E. coli* based on minimal reduction in viability and partial reduction (*P* > 0.05) in growth rates over time (Fig. 1). However, upon exposure to a moderate dose, there was a partial killing effect on *E. coli* after the first hour, followed by a slight increase in growth after the second and third hours (*P* > 0.05; Fig. 1A). The slight increases observed during the second and third hours indicate a ‘shock’ effect, which appeared to be short-lived based on apparent recovery after prolonged exposure. The mean bacterial cell recovery for *E. coli* O157:H7 at a high dose was low after the first hour (*P* > 0.05). Further reduction in bacterial survival continued on with longer exposure through the second and third hours (*P* < 0.05; Fig. 1A).

**Figure 1.**
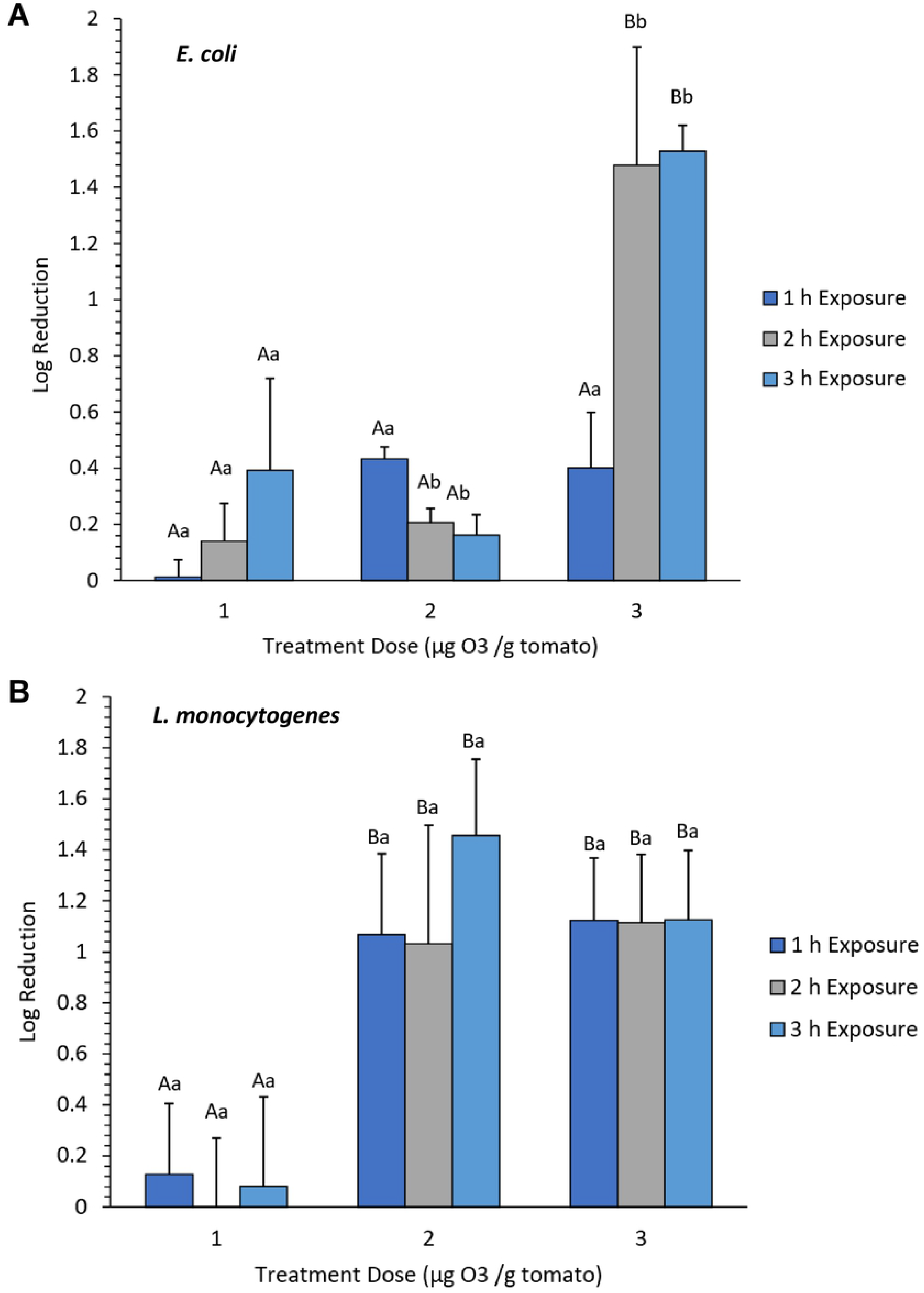
Growth reduction of *E. coli* **(A)** and *L. monocytogenes* **(B)** on non-host tomato treated with different doses of gaseous ozone (O_3_). Tomatoes harboring bacterial cells were treated with 1 μg, 2 μg, and 3 μg O_3_ per gram of ripe fruits. Capital letters indicate significant difference (P < 0.05) caused by O_3_ doses at the same time-point. Lowercase letters indicate significant difference (P < 0.05) caused by time of exposure under the same O_3_ dosage. (PowerPoint 106 KB)

Similar to the trends observed in *E. coli*, the effects of low dose (1 μg) and short duration (1 hr) exposure on the viability of *L. monocytogenes* on non-host tomato was very mild (Fig. 1B; *P* > 0.05). However, moderate and high doses caused a significant reduction in survival as indicated by the decline in bacterial counts even after short (1 hr) exposure. The effects of moderate and high doses continued through the second and third hours, although the magnitude during prolonged exposure appeared to be relatively mild (*P* < 0.05; Fig. 1B). In summary, the bacterial regrowth assays for *E. coli* and *L. monocytogenes* from the O_3_-treated fresh tomatoes across different doses and exposure time indicate that the impact of O_3_ xenobiosis was generally mild and appeared to have different levels of efficacy in the Gram-negative *E. coli* and Gram-positive *L. monocytogenes.* While the dose effects were similar between the two species of bacteria, the effect of exposure time appeared to vary. These trends appeared to indicate that the response mechanisms of the Gram-negative *E. coli* and Gram-positive *L. monocytogenes* are quite distinct.

Determining the optimal dose and exposure time that could induce effective killing of bacterial contaminants on the surface of a labile, non-host fresh produce such as tomato is a critical aspect of effective chemical intervention. Optimal dose and exposure time that maintain the overall physical and biochemical properties of the fresh produce are of prime importance. In the case of O_3_, a dose of 3 μg per gram of tomato fruits for up to 3 hr of treatment is the highest possible strength that can be applied without drastic impacts on sensory attributes and quality. Our earlier studies indicated that the highest efficacy of ClO_2_ xenobiosis requires a moderate dose and long exposure, which was not observed in the O_3_ assays for both *E. coli* and *L. monocytogenes* [33]. Therefore, based on the results of bacterial regrowth assays across dose and time combinations, O_3_ appeared to be less effective than ClO_2_ in reducing bacterial contamination on fresh tomatoes. O_3_ causes only mild killing effects on both *E. coli* and *L. monocytogenes* even at the highest dose and time exposure within the threshold level that maintains the overall sensory quality of tomato fruits. Compared to the overall effects of ClO_2_, given that only partial killing could be achieved at best with O_3_, we hypothesize that its utilization as a chemical agent for intervention has a higher likelihood of inducing adaptation and acclimation effects than ClO_2._

### Transcriptomic changes in response to O_3_ xenobiosis in *E. coli* and *L. monocytogenes*

We profiled the transcriptomes of *E. coli* and *L. monocytogenes* in the context of dose × time responses as a means to understand the nature of defenses and how those defenses are compromised when O_3_ xenobiosis triggered partial killing effects [30]. RPKM expression values across the temporal RNA-Seq datasets were normalized and transformed to Propensity Scores (PS). Pearson Correlation Coefficient (PCC) was further applied to PS in order to establish the significance of differentially expressed genes [30]. Based on the scatter plots of the global O_3_-response transcriptomes, similar patterns of transcriptional changes were evident over time during exposure of *E. coli* to low and moderate doses of O_3_ (Fig. 2A; Additional File 1: Fig. S). The responses triggered by low and moderate doses were characterized by the general downregulation of gene expression, which was particularly more pronounced during the second hour and continued through the third hour. On top of the general patterns of downregulation, there appears to be a certain subset genes that were upregulated during the second hour.

**Figure 2.**
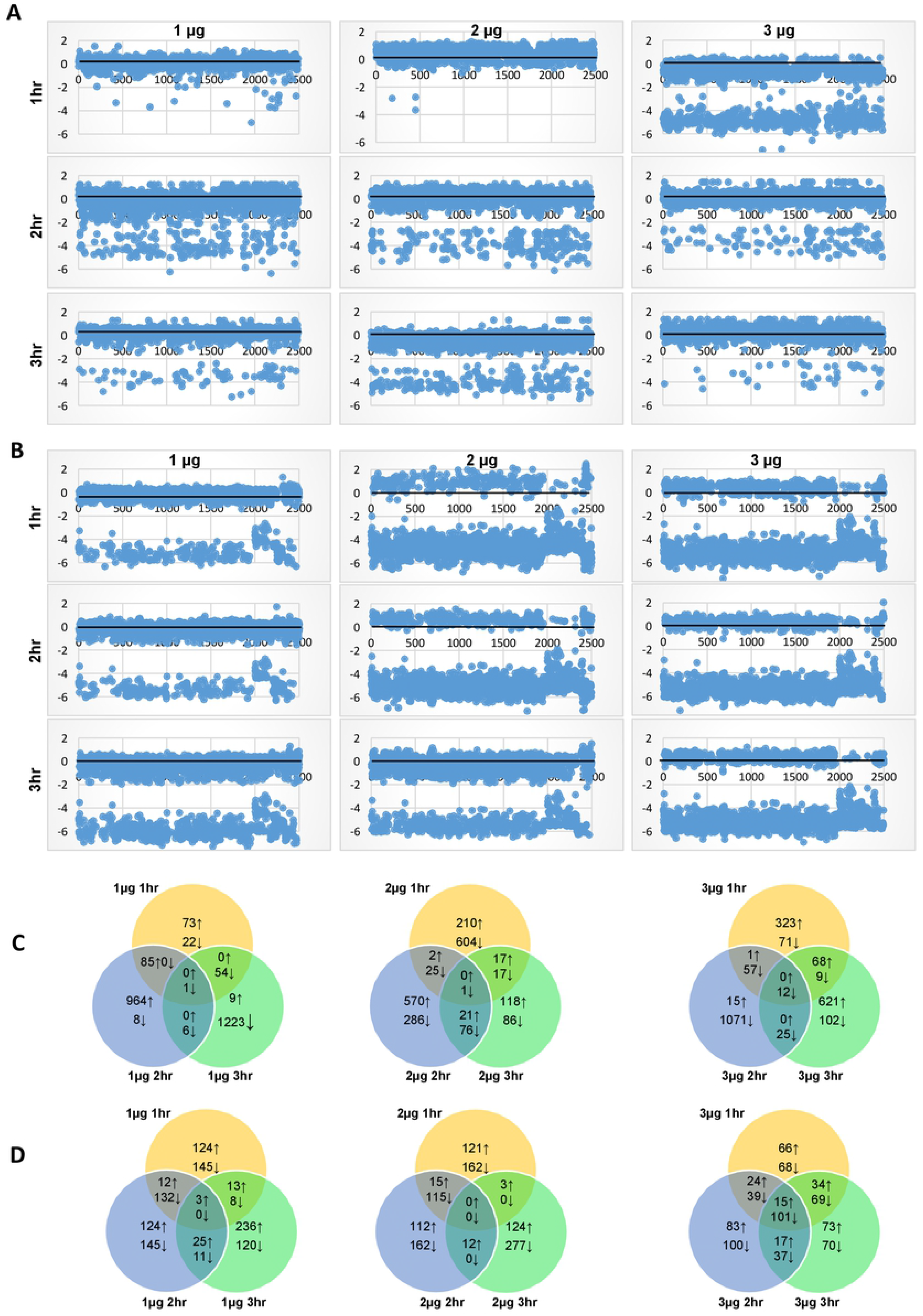
Dynamic changes in the *E. coli* O157:H7 and *L. monocytogenes* transcriptomes as an effect of different doses of O_3_ treatments. **(A and B)** Scatter plots of the Propensity Scores (PS) for each transcriptome library in *E. coli* O157:H7 **(A)** and *L. monocytogenes* **(B)**. **(C and D)** Total numbers of *E. coli* **(C)** and *L. monocytogenes* **(D)** genes that were differentially expressed in response to 1 μg, 2 μg, and 3 μg of O_3_ per gram of ripe tomato after 1 hr, 2 hr, and 3 hr exposure at each dose. Total numbers of upregulated (↑) and downregulated (↓) genes that were either treatment-specific or shared between treatments are displayed in the Venn diagrams. (PowerPoint 733 KB)

The global O_3_-response transcriptome of *E. coli* under a high dose is characterized by even more significant downregulation of gene expression, which was evident as early as during the first hour. The apparent perturbation appeared to suggest that higher doses are much more effective, although the xenobiotic effects are not quite evident during short duration (first hour) exposure as indicated by the growth reduction data (*P* > 0.05; Fig. 1A; Fig. 2A). Strikingly, as O_3_ treatment is prolonged through the third hour, the response transcriptome appeared to show indications of recovery from the initially perturbed state as indicated by the significant reversal of downregulated genes towards upregulation (Fig. 2A). These signs of recovery correlated with the results of the regrowth assay, suggesting that reduction in growth due to xenobiosis was substantially attenuated during the second and third hours (*P* < 0.05; Fig. 1A). These trends further suggest a gradual recovery of the surviving bacterial sub-populations, which are likely consequences of potential adaptation due to chronic effects.

In stark contrast, the global O_3_-response transcriptome of *L. monocytogenes* under a low dose appeared to be more perturbed as indicated by the widespread downregulation of gene expression during the entire period (first to third hours) of treatment (Fig. 2B). This trend seemed to be contradictory to the patterns in regrowth assay when reduction in bacterial regrowth was not very apparent (Fig. 1B). At moderate to high doses, the global transcriptomes were apparently perturbed, as indicated by the massive downregulation of gene expression throughout the entire three-hour duration of exposure. This trend was consistent with a significant reduction in bacterial viability (*P* < 0.05) as indicated by the regrowth assay (Fig. 1B).

The RNA-Seq data matrix revealed that the expression of a total of 2,488 genes in *E. coli* (48.5% of total protein-coding genes) and 1,801 genes in *L. monocytogenes* (63.5% of total protein-coding genes) were altered during the three-hour treatment period, as evident from different patterns of downregulation and upregulation across time (Fig. 2C and 2D; Additional file 2: Table S1). In *E. coli,* a total of 73, 964, and 9 genes were uniquely upregulated in response to low dose during the first, second and third hours, respectively. In *L. monocytogenes*, there were 22, 8 and 1,233 uniquely upregulated genes in response to low dose during the first, second and third hours, respectively. These temporal changes in gene expression indicate a gradual response to a low dose of O_3_ that appeared to peak with longer periods of exposure. The significant spikes in the number of upregulated genes during the third hour indicate that defenses are progressively induced with longer exposure, which is consistent with the overall patterns in the regrowth assay where log-reduction in bacterial growth was only mild even with a longer duration of exposure (Fig. 1C).

At a moderate dose, a total of 210, 570, and 118 genes in *E. coli* were uniquely upregulated during the first, second, and third hours, respectively. Moderate dose also caused the downregulation of 604, 286, and 86 genes in during the first, second, and third hours, respectively (Fig. 2C). These opposite patterns in the upregulation and downregulation of gene expression during prolonged exposure suggest that defense responses are gradually stabilized over time under a non-lethal dose of O_3_. Gradual stabilization of defense responses implies that perturbation is diminished or repaired over time under non-lethal dose, which is consistent with the resurgence of bacterial growth based on the higher proportions of viable cells recovered during the third hour of treatment as indicated by the regrowth assay (Fig. 1A).

During the first hour of exposure to a high dose, 392 genes were upregulated, and 194 genes were downregulated in *E. coli* (Fig. 2C). Notably, there was a drastic decline in the number of upregulated genes from 392 to only 16 during the second hour, which was accompanied by a spike in the total number of downregulated genes from 194 to 1,165 (Fig. 2C). During the third hour, a total of 689 genes were upregulated, and 148 genes were downregulated (Fig. 2C). The large proportion of downregulated genes during the second hour suggests that severe perturbation hence killing effects have occurred, consistent with the log-reduction data (Fig. 1A). However, the spike in the number of upregulated genes during the third hour suggests significant recovery, likely as a result of acclimation during prolonged exposure (Fig. 1A).

Exposure of *L. monocytogenes* to a low dose caused the upregulation of 124, 124, and 236 genes during the first, second, and third hours of treatment, respectively, and downregulation of 145, 145, and 120 genes during the first, second, and third hours, respectively (Fig. 1D). These changes imply that at low dosage, defense responses in *L. monocytogenes* are not fully active unless the bacterial population is subjected to prolonged exposure. Under moderate dose, which caused significant reduction in bacterial regrowth (Fig. 1B), a total of 121, 112, and 124 genes were uniquely upregulated, during the first, second, and third hours, respectively, while 162, 162, and 277 genes were uniquely downregulated during the first, second, and third hours, respectively. These trends in gene activation and repression across time are indicative of attenuated defense response, particularly during longer exposure (Fig. 2D). Further increase to a high dose caused the upregulation of 66, 83, and 73 genes during the first, second, and third hours, respectively, and downregulation of 68, 100, and 70 genes during the first, second, and third hours, respectively (Fig. 2D). During exposure of *L. monocytogenes* to a high dose, much larger number of differentially expressed genes overlapped at during short, medium and longer duration, which are suggestive of potential adaptation.

### Chronic exposure causes a new burst of defenses

Exposure of *E. coli* to a low dose of O_3_ was accompanied by upregulation of 815 genes during the second hour, while the same subset of genes drastically shifted to downregulation during the third hour (Additional file 2: Table S1). Under a moderate dose, a subset of 346 genes that were downregulated during the first hour shifted to upregulation during the second hour. These drastic shifts in gene expression indicate that a new burst of defense responses may have been triggered likely due to the intense selection pressure associated with longer exposure even at moderate dose. We also observed that a subset of 423 genes that were downregulated during the second hour at high dose drastically shifted to upregulation during the third hour. This suggests that potential acclimation of the surviving sub-populations during prolonged exposure may have likely occurred.

Under low dose, few genes in *L. monocytogenes* that were upregulated during the second and third hours shifted to downregulation with longer exposure to moderate (2 μg) and high (3 μg) doses. This subset of genes is enriched with regulatory functions that are important for defense mechanisms, including the Rrf2 family protein gene (LMOf2365_2331), NifU family protein (LMOf2365_2371), FUR family transcriptional regulator (LMOf2365_1986), and GlnR family transcriptional regulator (LMOf2365_1316) (Additional file 2: Table S1).

In both *E. coli* and *L. monocytogenes,* the peculiar gene expression signatures associated with dose x time response were enriched with functions associated with pathogenicity, response to stress, regulation of cell division, cell motility, amino acid and protein metabolism, transcription and RNA processing, transport, carbohydrate metabolism, nucleotide metabolism, and genetic recombination (Fig. 3; Additional file 2: Table S1). In *E. coli*, a large proportion of genes associated with pathogenesis and stress response were progressively downregulated from the first to the second hour under low dose. A significant proportion of these genes shifted to upregulation during the third hour (Fig. 3A). With a further increase in dose to moderate, genes associated with pathogenesis and stress response were either upregulated or downregulated across time (Fig. 3A). However, under a high dose, a much larger proportion of pathogenesis and defense response genes were significantly downregulated during the first and third hours but upregulated during the second hour. These trends are suggestive of a new burst of expression of defense-associated genes, an event that is likely independent of the responses that occurred during the first hour (Fig. 3A). In contrast, genes associated with pathogenicity, stress response, cell division, and cell motility were upregulated in *L. monocytogenes* during exposure to low, moderate, and high doses across time (Fig. 3B).

**Figure 3.**
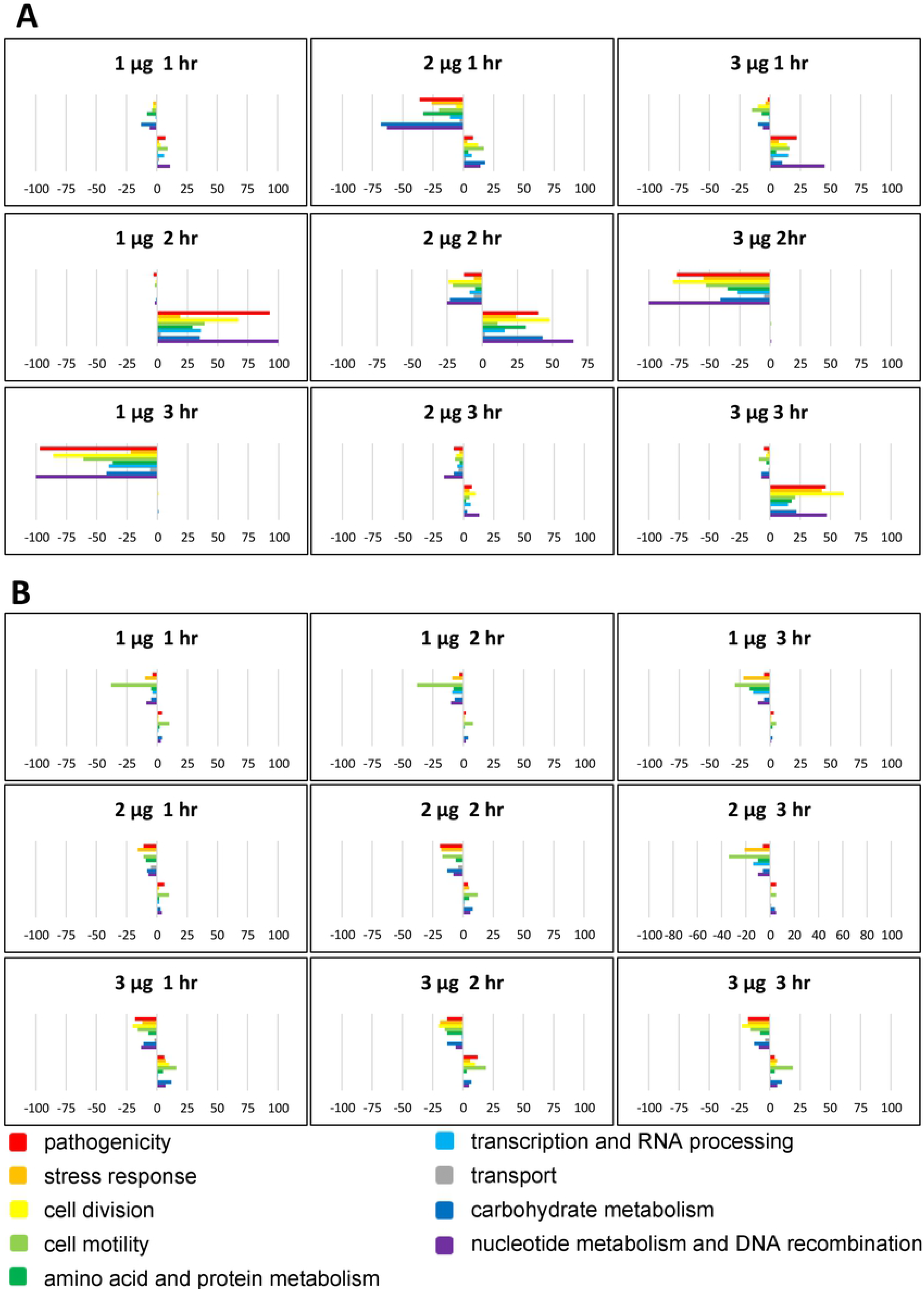
Functional categories of genes in *E. coli* O157:H7 **(A)** and *L. monocytogenes* **(B)** that were differentially expressed at a different duration of exposure (1 hr, 2 hr, 3 hr) to O_3_. Colored bars represent the numbers of differentially expressed genes assigned to each functional category. Positive bars denote the number of upregulated genes. Negative bars denote the number of downregulated genes. Genes assigned to ‘other category’ and ‘unknown’ are not included in this figure. (PowerPoint 111 KB)

Two-way hierarchical clustering of Propensity Scores (PS) and RPKM values revealed 20 distinct clades representing different patterns of co-expression across the *E. coli* and *L. monocytogenes* transcriptomes (Fig. 4; Additional file 2: Table S1). In *E. coli*, clustering of PS according to dose x time regimes showed that expression signatures in 1 μg O_3_ × 1 hr treatment regime were more similar to the signatures of 3 μg O_3_ × 1 hr treatment regime, while the expression signatures of 1 μg O_3_ × 2 hr regime clustered with the signatures of 3 μg O_3_ × 3 hr regime (Fig. 4A). The expression signatures based on RPKM data indicate that the profiles of 1μg O_3_ × 1 hr and 1 μg O_3_ × 2 hr treatment regimes formed a single clade, while the profiles of 1 μg O_3_ × 3 hr and 3 μg O_3_ × 3 hr treatment regimes formed a separate clade (Fig. 4A). In *L. monocytogenes*, the gene expression signatures of 1 μg O_3_ × 1 hr, 1 μg O_3_ × 2hr, 1 μg O_3_ × 3 hr, and 2 μg O_3_ × 3 hr treatment regimes were more similar, characterized by more significant downregulation. Expression signatures of the other treatment regimes formed a separate clade characterized by significant upregulation (Fig. 4B). These trends indicate that defense responses were attenuated at a low dosage but much enhanced at a higher dosage.

**Figure 4.**
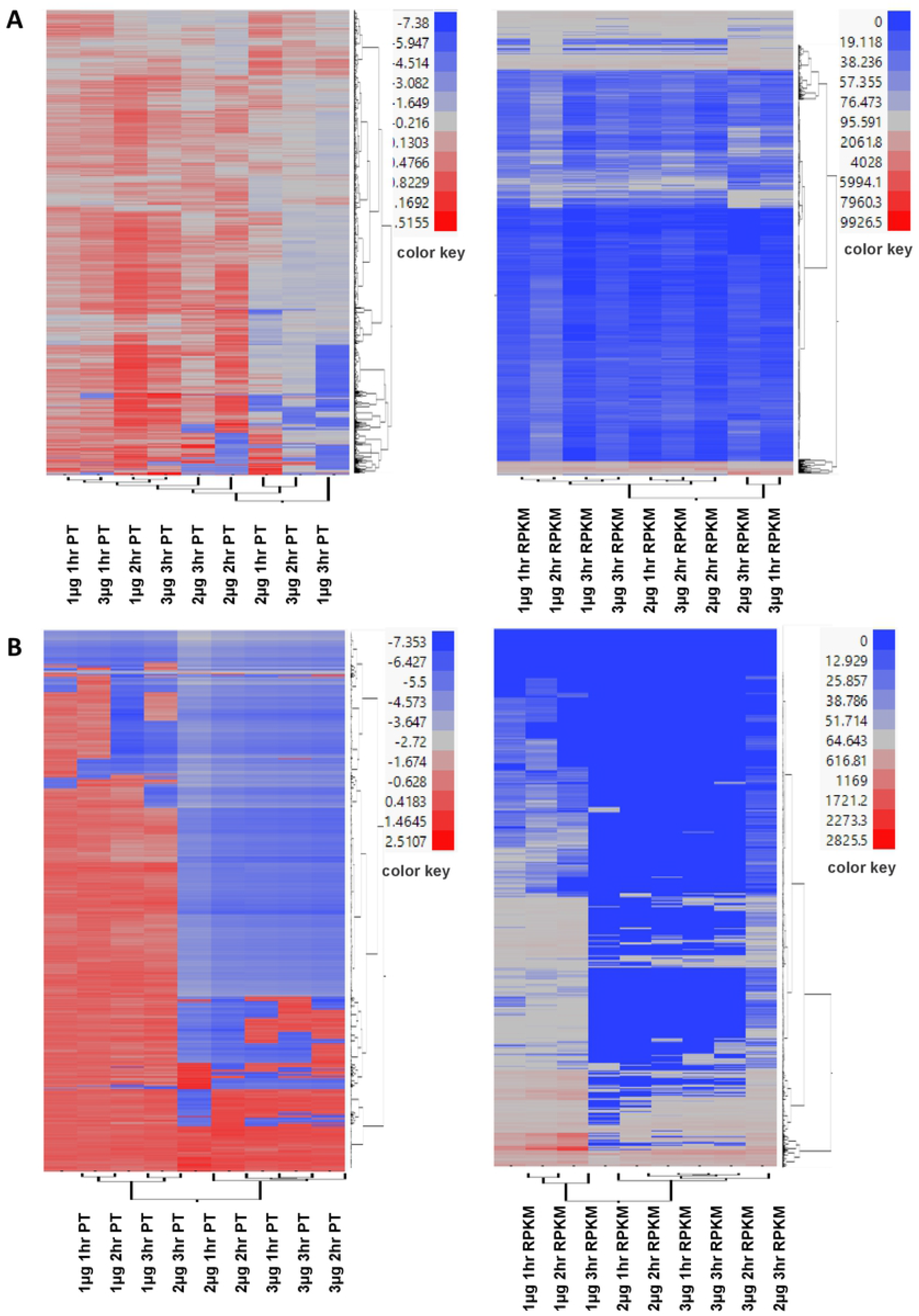
Two-way hierarchical clustering of differentially expressed *E. coli* **(A)** and *L. monocytogenes* **(B)** genes. Ward’s Hierarchical Clustering was performed to analyze the Propensity Scores (PS) (left) and RPKM values (right) of bacterial genes in response to O_3_ treatments after 1 hr, 2 hr and 3 hr exposure. The number of clusters was set at 20 with color-coding, as shown in detail Additional file 2: Table S1. Red indicates high expression and green indicates low expression. (PowerPoint 163 KB)

### Effects of O_3_ on genes associated with pathogenicity, stress response, and defenses

T3SS represents an important class of genes involved in bacterial pathogenicity, encoding multi-protein complex channels that inject effectors to promote bacterial attachment to the host [34]. The transcriptomes of *E. coli* revealed a total of 29 T3SS-encoding genes with altered expression, particularly in response to high dose (Fig. 5A; Additional file 2: Table S1). Of these genes, 17 were upregulated in the 3 μg O_3_ × 1 hr treatment regime, and 10 were upregulated in the 3 μg O_3_ × 3 hr treatment regime. An additional 21 T3SS-encoding genes were downregulated in the 3 μg O_3_ × 2 hr treatment regime (Fig. 5A; Additional file 2: Table S1). Differential expression of these T3SS genes indicates that xenobiotic effects at high dose compromised pathogenicity, particularly with moderately extended time up to 2 hours. During the third hour, there is an indication of a rebound of pathogenicity, likely as a consequence of prolonged exposure.

**Figure 5.**
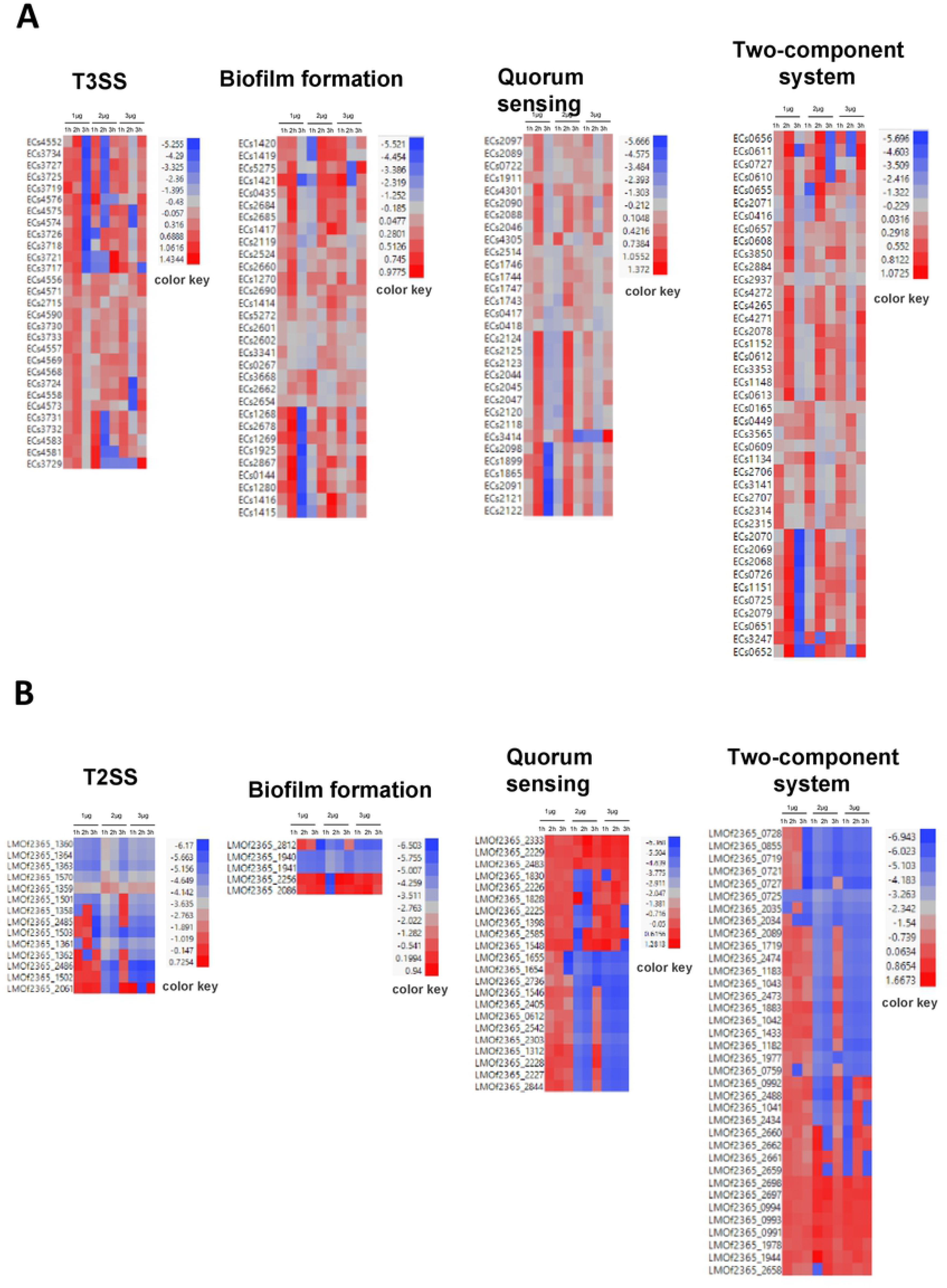
Heat map of differentially expressed *E. coli* **(A)** and *L. monocytogenes* **(B)** genes associated with pathogenesis and stress response. PS of genes in response to O_3_ treatments after 1 hr, 2 hr and 3 hr were plotted. Red indicates high expression and green indicates low expression. (PowerPoint 122 KB)

In *L. monocytogenes*, expression of 14 genes that belong to the general class of T2SS-pathogenesis proteins were affected by exposure to O_3_. Of these, five (5) genes encoding ATP-binding protein (LMOf2365_1360), comEC/Rec2 family protein (LMOf2365_1501), general secretion pathway protein E (LMOf2365_1364), and general secretion pathway protein F (LMOf2365_1363) were downregulated in 1 μg O_3_ × 1 hr treatment regime, and another helicase-encoding gene (LMOf2365_2486) was stably downregulated throughout the three-hour period of O_3_ treatment (Fig. 5B; Additional file 2: Table S1).

A total 18 genes involved in biofilm formation were affected by O_3_ treatments in *E. coli* (Fig. 5A; Additional file 2: Table S1). Under low dose, several of these biofilm-associated genes were upregulated during the first and second hours, and then subsequently downregulated during the third hour. At a higher dose, another subset of biofilm-associated genes that were initially downregulated during the first hour were subsequently upregulated through the second and third hours. With further increase to a high dose, another subset of biofilm-associated genes was upregulated during the first and third hours but downregulated during the second hour. These expression signatures imply that the process of biofilm formation was attenuated during brief exposure to O_3_ regardless of the dosage, but the same process appeared to be enhanced with longer exposure to much higher doses with longer exposure. In contrast to the large effects of O_3_ on biofilm-associated genes in *E. coli*, only four (4) biofilm-associated genes were affected in *L. monocytogenes*, including those encoding diguanylate cyclase (LMOf2365_1940, LMOf2365_1941), which were downregulated during the first hour at low dose (Fig. 5B; Additional file 2: Table S1).

As a critical component of cell-to-cell communication in bacteria, quorum sensing facilitates the monitoring of cell population density, detection of xenobiotic molecules, and translation of extracellular signals to intercellular processes and downstream gene expression [35]. The response transcriptomes of *E. coli* and *L. monocytogenes* included 31 and 15 quorum sensing-associated genes, respectively, that were significantly affected by O_3_ (Fig. 5; Additional file 2: Table S1). In *E. coli*, these genes were downregulated during short (1 hr) and long (3 hr) duration exposure to low dose. At a moderate dose, these genes were downregulated during the first and third hours but upregulated during the second hour. With further increase to a high dose, these genes were upregulated during the first hour, downregulated during the second hour, and subsequently dramatically upregulated during the third hour. In *L. monocytogenes*, two (2) quorum sensing genes were upregulated during the first and second hour at a low dose, and another gene was downregulated during the third hour. However, at moderate and high doses, quorum sensing genes exhibited different patterns of upregulation or downregulation across time, suggesting that a comprehensive defense system is triggered *L. monocytogenes* under higher strength of O_3_ xenobiosis.

The two-component system plays an important role in bacterial responses to environmental changes, which is critical in maintaining pathogenicity and fitness under adverse conditions [36]. The transcriptome data revealed a total of 41 and 21 genes associated two-component systems that were affected by O_3_ in *E. coli* and *L. monocytogenes,* respectively (Fig. 5; Additional file 2: Table S1). Of the 41 affected genes in *E. coli*, 24 were significantly upregulated at low dose during the second hour, while 21 genes were downregulated during the third hour. At higher dose, several two-component system-associated genes were upregulated during the first and third hours but were attenuated during the second hour. As mentioned in the previous section, transient upregulation of a large number of genes at high dose with longer exposure are likely indicators of acclimation possibly due to chronic effects (Fig. 1). In *L. monocytogenes*, most of the genes associated with the two-component system were downregulated, with only a few being upregulated, particularly under high dose, suggesting that O_3_ has a more significant impact in suppressing the two-component system in *L. monocytogenes* than *E. coli*.

In addition to the effects on T3SS/T2SS, biofilm, quorum-sensing, and two-component associated genes, O_3_ also had significant effects on other classes of stress-related genes in both *E. coli* and *L. monocytogenes*. For instance, in *E. coli*, several genes encoding SOS response proteins, thiol:disulfide interchange protein, putative tellurium resistance protein, and chemotaxis-associated proteins were affected at different time-points (Additional file 2: Table S1). In addition, six heat shock protein genes (HSPs) were upregulated during the third hour under low dose but downregulated during the first hour under moderate dose. With a further increase to a high dose, genes that function in the regulation of energy metabolism [37] as well as phosphotransferase system HPr enzyme (ECs4354) were downregulated during the second hour but upregulated during the third hour.

In *L. monocytogenes,* genes associated with stress response were under high dose, including an amidophosphoribosyltransferase (LMOf2365_1793), 2-oxoisovalerate dehydrogenase E1 subunit beta (LMOf2365_1390), transketolase (LMOf2365_2640), and ABC transporter ATP-binding protein/permease (LMOf2365_2732). Other types of stress-related genes such as ribose-5-phosphate isomerase B (LMOf2365_2654), heat shock protein GrpE (LMOf2365_1493), and heat-inducible transcription repressor (LMOf2365_1494) were upregulated. Few genes involved in antibiotic biosynthesis were also affected by O_3_ in both *E. coli* and *L. monocytogenes*, indicating that O_3_ elicit responses similar to the defense mechanisms against antibiosis.

Certain genes involved in the regulation of bacterial transcription such as *sigma-E factor* are key players in biofilm formation and pathogenicity [38–40]. The transcriptome data revealed that O_3_ had an effect on the expression of certain transcriptional regulatory proteins associated with biofilm formation and pathogenicity in both *E. coli* and *L. monocytogenes.* For example, in *E. coli*, the *sigma-E regulatory protein* gene (ECs3436) was upregulated during the first hour and then subsequently downregulated during the second hour under moderate dose (Additional file 2: Table S1). Potential suppression of biofilm-associated genes appeared to be supported by concomitant downregulation of translational activities as suggested by the downregulation of 12 ribosome-associated genes. In *L. monocytogenes*, the *transcription termination factor Rho* (LMOf2365_2523) was downregulated during the entire period (first to third hour) under high dose.

### Transcriptional networks associated with bacterial responses to O_3_

We selected the genes that were most significantly affected by O_3_ in a dose x time dependent manner in both *E. coli* and *L. monocytogenes.* These included the genes with the lowest and highest PS across the three doses used in the experiments. A cut-off of 0.9999 was further applied to filter out both positively and negatively correlated changes in gene expression. At this stringency of filtration, 1,588, 1,276 and 1,405 genes in *E. coli* represent the most statistically significant changes in expression during the first, second, and third hours, respectively. Similarly, 1,221, 1,312 and 1,119 genes in *L. monocytogenes* represent the most statistically significant changes during the first, second and third hours, respectively. These groups of genes were used to model the transcriptional co-expression networks to understand the effects of O_3_ xenobiosis on the global response mechanisms of each bacterial species (Additional file 3: Table S2). Pearson Correlation Coefficients (PCC) identified 674, 781, and 934 genes that were differentially expressed in *E. coli* under low (1 μg), moderate (2 μg), and high (3 μg) doses_3_, respectively (Additional file 2: Table S1; Additional file 3: Table S2). Similarly, PCC identified 876, 591, and 736 genes that were differentially expressed in *L. monocytogenes* under low, moderate and high doses, respectively (Additional file 2: Table S1; Additional file 3: Table S2).

Under low dose, the transcriptional network of *E. coli* during the first hour is characterized by few small clusters of co-expressed genes at the center of the global network (Fig. 6A). During the second hour, large clusters of genes appeared to be coordinately regulated in either positive (co-upregulation) or negative (co-downregulation) direction without an apparent connection to a central hub or core regulator. During the third hour, larger clusters of co-expressed genes started forming a discernible connection to a central hub or core regulator. Under moderate and high doses, the networks included few small reorganized secondary clusters during the entire three hours of exposure (Fig. 6A). This trend appears to suggest that high doses possibly led to acclimation, based on the reduced magnitude of gene expression changes relevant to the defense. These results indicate that *E. coli* responds to low and high doses in very different ways, similar to the dose-dependent responses reported earlier for ClO_2_ [33].

**Figure 6.**
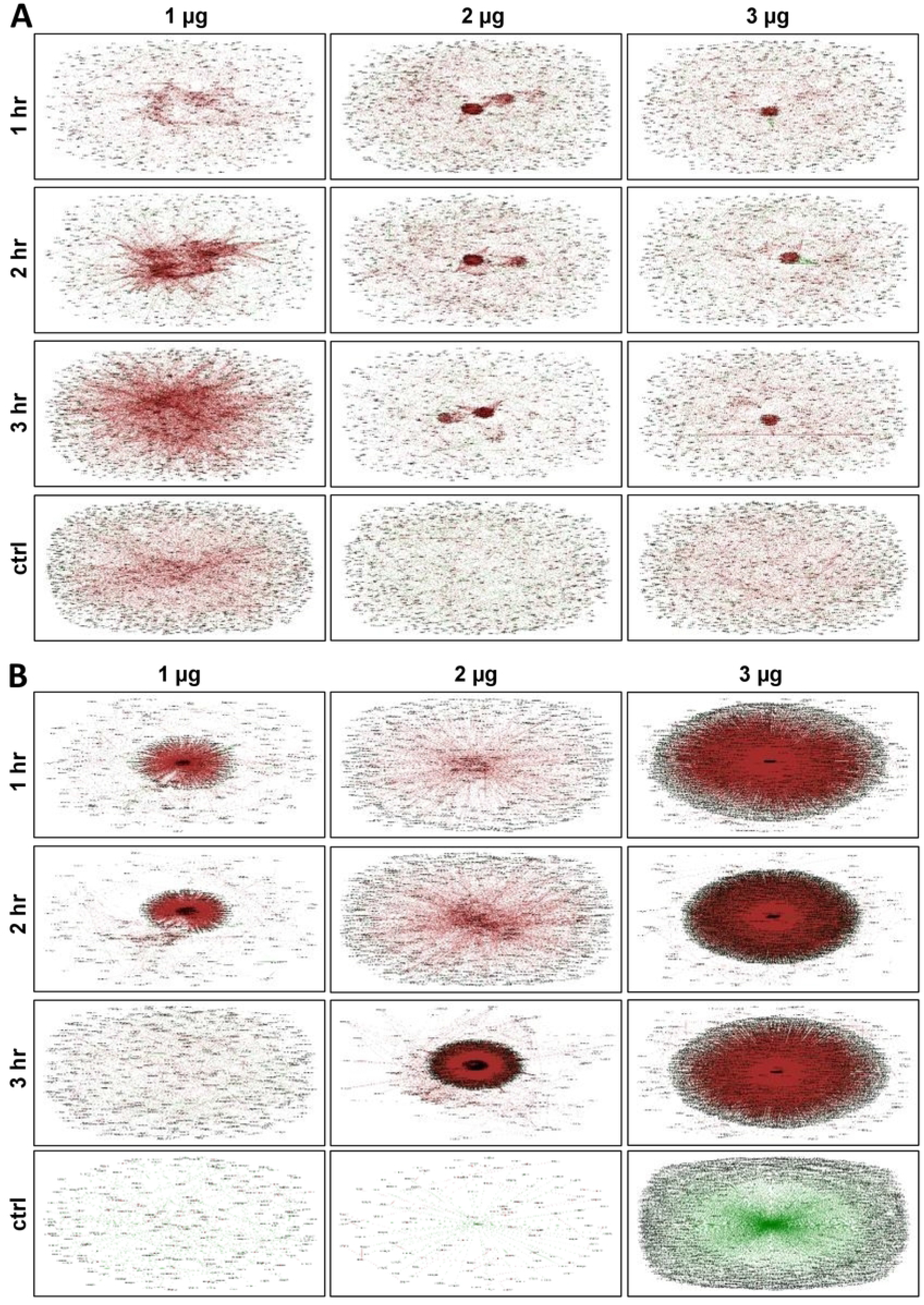
Models of transcriptional co-expression networks constructed for *E. coli* **(A)** and *L. monocytogenes* **(B),** based on co-expression under control (untreated), and 1 hr, 2 hr, and 3 hr exposure to different doses of O_3_. Each node represents a gene and each line denotes the expression correlation between the two nodes. Green node denotes upregulated genes; blue node denotes downregulated genes; brown line denotes positively correlated genes by Pearson Correlation Coefficient (PCC); green line denotes negatively correlated genes by PCC. (PowerPoint 2,172 KB)

The O_3_-response networks of *L. monocytogenes* appeared to be quite distinct from those of *E. coli*, supporting our hypothesis that *L. monocytogenes* is more sensitive to O_3_ as indicated by the regrowth assays (Fig. 1A). Under low dose O_3_, a small cluster of co-expressed genes was evident at the center of the global network during the first and second hours but not during the third hour (Fig. 6B). With further increase to a moderate dose, small clusters began to form at the center of the global network particularly during the first and second hour, which appeared to have resulted to a much larger cluster during the third hour (Fig. 6B). Under high dose, large clusters were evident during the entire three-hour duration of exposure (Fig. 6B).

## Discussion

### Acclimation of bacterial pathogens to non-host intermediate vectors

Gram-positive bacteria have cell walls made of a thick layer of peptidoglycan, while the cell walls of Gram-negative bacteria are composed of a thin layer of peptidoglycan and an outer membrane that is absent in Gram-positive bacteria. Gram-negative and Gram-positive bacteria have employed different molecular strategies to cope with the environmental changes and to interact with their hosts [41]. The Gram-negative *E. coli* and Gram-positive *L. monocytogenes* are among the most dangerous bacterial pathogens notorious for their ubiquitous occurrence across a broad range of environments. Their persistent nature and strong virulence have been attributed to their toxin production capacities and low infectious doses, causing high mortality rates in both humans and animals [3, 7, 42–46]. For instance, illnesses caused by the foodborne Shiga toxin-producing *E. coli* O157 (STEC) can be life-threatening, with a very low dose (20 and 700 cells) in contaminated fresh produce capable of causing major outbreaks [47]. In recent times, *E. coli* serotype O157:H7 has been the major cause of outbreaks by contaminating fresh vegetables during the post-harvest processing pipeline [7]. *E. coli* O157:H7 is flexible in terms of its adaptability to extreme fluctuations in the environment, due in part to its short life cycle and highly efficient genetic regulatory machineries that confer highly flexible defense systems [48, 49]. Harsh environmental conditions are largely responsible for triggering viable but non-culturable (VBNC) populations of bacteria, which provide an effective inoculum when resuscitated under the right environmental conditions [50]. The highly infectious *L. monocytogenes* cause high mortality rate regardless of antibiotic treatments [46]. Listeriosis disease caused by *L. monocytogenes* often leads to rare complications that are highly threatening to human health [51]. As various types of chemical intervention strategies are continuously developed to combat the potent and recurring infectious agents such as *E. coli* and *L. monocytogenes*, potential contributions of such intervention strategies to the evolution of resilience among the newly emerged isolates are often overlooked. Acclimation and adaptation to strong selection pressures (supra-optimal effects) triggered by xenobiotic agents must be well understood for more strategic implementation of combinatorial approaches to intervention.

With the increasing social and economic burdens caused by bacterial anti-microbial resistance (AMR), the World Health Organization (WHO) reported that significant gaps continue to emerge during the development of a successful anti-microbial stewardship program [52]. Given its medical, environmental, and industrial importance, a comprehensive understanding of bacterial response to chemical intervention is essential in combatting bacterial pathogens and their potential global socio-economic impacts. Most studies that investigated how bacterial pathogens respond to chemical hygiene practices mainly focused on developing new strategies to diminish contamination [21]. Our previous studies demonstrated that the efficacy of growth reducing action caused by another chemical intervention agent ClO_2_ against *E. coli* on on-host tomato was dependent on dose × time effects [33]. Previous studies have unraveled the cellular and molecular mechanisms of β-lactam antibiotic-induced resistance in *E. coli* [53]. Despite this advance, little is known about the nature of bacterial defense responses triggered by supra-optimal dose × time effects. Effective, and sustainable treatment strategies that reduce contamination of fruits and vegetables have been difficult to achieve, largely due to poor understanding of the molecular genetic mechanisms associated with xenobiotic effects of chemical intervention agents. Fine-tuned strategies that balance the dose x time effects with the need to preserve the sensory and biochemical properties of the fresh produce are critical in preventing the potential supra-optimal effects of xenobiotic agents to bacterial acclimation, adaptation, and even mutation.

Our previous studies on the effects of ClO_2_ to *E. coli* on non-host tomato environment revealed that supra-optimal exposure time, even at an effective dose, could lead to a second burst of independent defense responses [33]. In particular, the activation of pathogenicity and stress-adaptive genes response with prolonged exposure to a high dose (10 μg) of ClO_2_ points to the probable occurrence of adaptation and selection, leading to resistance. However, we observed a new burst of defense responses triggered by a moderate dose of O_3_ (2 μg) in *E. coli* at the second hour of exposure (Fig. 2), an indication low dosage of O_3_ potentially triggers acclimation and adaptation.

### Xenobiotic effects of O_3_ against *E. coli* and *L. monocytogenes*

The potential of gaseous O_3_ as a xenobiotic agent for pathogen intervention on foods has been demonstrated [16-18, 54, 55]. In this study, we revealed that O_3_ is capable of reducing the level of viable *E. coli* and *L. monocytogenes* inocula on tomato fruits as a non-host vector for transmission to humans [8]. The transcriptional regulatory networks of both *E. coli* and *L. monocytogenes* under various conditions have been studied previously [56–60]. We investigated the transcriptional regulatory networks of *E. coli* and *L. monocytogenes* surviving on its non-host tomato after exposure to low (1 μg per gram of fruit), moderate (2 μg per gram of fruit) and high (3 μg per gram of fruit) doses of O_3_. Our results indicated that *E. coli* and *L. monocytogenes* respond to O_3_ exposure in dosage × time-dependent manner (Fig. 1, 2 and 5). High dose appeared to have the highest potential to trigger adaptation in *E. coli* but not in *L. monocytogenes,* suggesting that *E. coli* has higher basal resistance than *L. monocytogenes* (Fig. 3). In our previous studies, we also reported that another xenobiotic intervention agent ClO_2_ induced potential acclimation and adaptation in *E. coli* surviving on fresh tomato [33]. Taking together, our findings suggest that when optimizing xenobiotic intervention procedures, potential bacterial adaptation needs to be taken into consideration while balancing the dose × time effects on the physical and biochemical properties of the fresh produce being subjected to such treatments.

### The nature of *E. coli* and *L. monocytogenes* response mechanisms to O_3_ xenobiosis

We revealed that O_3_ caused changes in the expression of genes associated with pathogenicity, stress response, cell motility, transcriptional regulation, primary metabolism, and transport in both *E. coli* and *L. monocytogenes* (Fig. 3). Pathogenicity genes involved in T3SS system (in *E. coli*), T2SS system (in *L. monocytogenes*), biofilm formation, quorum sensing, and two-component system were triggered during exposure to O_3_ (Fig. 5; Additional file 2: Table S1). When *E. coli* was exposed to a high dose of O_3_, upregulation of genes associated with those functions occurred largely during the short-term (1 hr) exposure, but such pattern of gene activity was not observed during prolonged exposure. In our previous study, we also observed changes in expression of T3SS system, biofilm formation, quorum sensing, and two-component system genes during exposure to ClO_2_ in *E. coli* [33].

T3SS genes involved in virulence (ECs4590, ECs3730, ECs3731, ECs3732, ECs3733, ECs3726, ECs3725, ECs3724 and ECs3721) were upregulated during short-term (1 hr) exposure to a high dose but downregulated with prolonged exposure up to 2 hr. Suppression of defense response was likely due to a significant reduction in metabolic activity, concurrent with partial or complete arrest of cell division [61] (Fig. 1A). The substantial similarities in the transcriptional changes in *E. coli* during moderate (2 hr) and longer (3 hr) exposure times but not with shorter (1 hr) exposure time indicated that longer exposure could potentially cause a selection pressure that could trigger acclimation and adaptation. The apparent exposure time-dependence of gene expression in *E. coli* under a high dose suggested that toxicity effects, as well as genetic mechanisms, might be different at various periods during xenobiosis. In contrast, these changes were not observed in *L. monocytogenes*, indicating adaptation is not likely induced in the Gram-positive bacteria as implied by the nature of its transcriptional networks (Fig. 3 and 6).

### Acclimation of *E. coli* and *L. monocytogenes* due to chronic effects

Our previous study showed significant changes in transcriptional regulatory networks in *E. coli* in response to ClO_2_ treatment as indicators of either defenses or acclimation [33]. In this study, systematic reconstruction of the transcriptional regulatory networks across different dosages of O_3_ (*i.e*., 1 μg, 2 μg, 3 μg) showed that gene modules associated with pathogenicity, stress response, transcriptional regulation, and transport processes play important roles in defense against O_3_ xenobiosis (Additional file 3: Table S2). In *E. coli*, prophage induction is often coupled with enhanced virulence and increased tolerance to harsh environmental stresses [62, 63]. The *E. coli* transcriptome data revealed a total of 196 prophage-associated genes to be differentially expressed across different doses and exposure times (Additional file 2: Table S1), indicating that prophage induction plays a critical role in the responses of *E. coli* to O_3_-mediated xenobiosis.

Current thinking supports that exposure to environmental stresses could stimulate mechanisms that enhance bacterial survival across different host or non-host environments, *i.e.,* stress priming effects [23, 24]. In *E. coli*, it is known that stress could induce acclimation, adaptation, selection, or even rare mutation events. Environmentally induced changes in fitness could lead to selection and population shift that build a novel inoculum with newly acquired tolerance to different modes of intervention. One example is that *E. coli* O157 is more resistant to acid once it is primed by heat treatment [25]. Cross-protection against other stresses induced by salt for example, has also been reported in *L. monocytogenes* in a temperature-dependent manner [64]. Whether O_3_ causes mutagenic effects and subsequently contributes to cross-protection mechanisms in *E. coli* or *L. monocytogenes* is unknown. From a food safety standpoint, introducing combinations of relatively mild chemical treatments (*i.e.,* optimal cocktail) might be an attractive alternative to effectively control bacterial pathogens without promoting adaptation or mutation, which are the main causes of perennial outbreaks.

Studies have shown that various types of stresses induce changes in gene expression in *E. coli* in both pure culture and non-host environments, such as fresh lettuce [48, 65]. In this study, we examined the dynamics of transcriptional co-expression networks of *E. coli* and *L. monocytogenes* in line with their responses to different doses of O_3_ on non-host tomato surface, which may serve as a pre-exposure to another stressor causing either cross-protection or cross-vulnerability [24]. We found that the genetic network configurations of *E. coli* and *L. monocytogenes* are very flexible under different doses of O_3_ over short or longer duration of exposure. We also characterized the transcriptional changes in *E. coli* and *L. monocytogenes* growing in pure culture and on tomato surface, providing great reference transcriptomes on these pathogens growing on various substrates (Additional file 2: Table S1).

We previously reported that the transcriptional regulatory network of *E. coli* in response to a low dose of ClO_2_ is controlled by a *putative endopeptidase* (ECs2739) as a central hub, likely through its functions associated with stress signaling, antibiotic binding and recognition, bacteriophage activity, and morphology determination [66, 67]. In this study, no putative central hub or core regulator was apparent for the O_3_ response networks of both *E. coli* and *L. monocytogenes*. However, it is apparent that the responses and associated mechanisms triggered by O_3_ xenobiosis are distinct from the responses triggered by ClO_2_. The *putative endopeptidase* gene that serves as the central hub in the ClO_2_ networks of *E. coli* was downregulated by 1 μg O_3_ at 3 hr and 3 μg O_3_ at 2 hr, but upregulated under 1 μg and 2 μg O_3_ at 2 hr, and 3 μg O_3_ at 3 hr (Additional file 2: Table S1). Such response mimics the typical profile of biological invasion that often involves the degradation of foreign proteins by enhanced endopeptidase activities [68, 69].

The current study illustrates the power of transcriptome profiling for understanding the genetic networks involved in the responses of pathogenic bacteria (*E. coli*, *L. monocytogenes*) to sub-optimal, optimal, or supra-optimal effects of a potential xenobiotic agent (O_3_) used for intervention in food processing. We have established a platform to investigate the molecular mechanisms underlying pathogen interaction with intervention chemicals, providing a baseline for optimizing dose × time dynamics for maximal efficacy [33]. The differentially expressed genes could serve as targets in both *E. coli* and *L. monocytogenes* for future development of novel strategies for controlling foodborne pathogens, including the use of new chemical and bio-control agents, *i.e.*, non-toxic and non-pathogenic biocontrol bacterial strains or phage can be considered. In addition, means for tricking the signal transduction pathways associated with defense response to various chemicals could be an alternative strategy to re-wire the bacterial genetic networks, thereby reducing selective pressure and avoiding the emergence of chemical-tolerant inocula. The information generated in this study also provides an important resource for further research in food safety and foodborne pathogen epidemiology.

### Conclusions

The present study provides a proof-of-concept on the potential xenobiotic effects of O_3_ to *E. coli* but not in *L. monocytogenes* and the importance of dose × time dynamics for optimal intervention. The paradigm of this study could be applied to evaluate the impacts of different intervention strategies in the food industry to eliminate bacterial pathogens surviving in fresh produce while minimizing the negative consequences on selection and adaptation.

### Data deposition

RNA-Seq data were deposited in the National Center for Biotechnology Information (NCBI) Sequence Read Archive (SRA) collection under the accession number SRR8468286-9.

## List of abbreviations

AMR: antimicrobial resistance
ClO_2_: chlorine dioxide
CT-SMAC: MacConkey Sorbital Agar supplemented with Cefixime and Tellurite
*E. coli*: *Escherichia coli*
hr: hour(s)
*HSP*: *heat shock protein*
*L. monocytogenes*: Listeria *monocytogenes*
NaB: sodium benzoate
NaClO: sodium hypochlorite
NaClO_2_: sodium chlorite
NCBI: National Center for Biotechnology Information
O_3_: ozone
PAA: peracetic acid
PCC: Pearson Correlation Coefficient
PS: propensity score
RNA-Seq: RNA-Sequencing
RPKM: Reads Per Kilobase of transcript, per Million mapped reads
SRA: Sequence Read Archive
STEC: Shiga toxin-producing *Escherichia coli*
T2SS: type II secretion system
T3SS: type III secretion system
TSA: tryptic soy agar
USDA: the U.S. Department of Agriculture
VBNC: viable but non-culturable
WHO: World Health Organization.

## Ethics approval and consent to participate

Not applicable

## Consent for publication

Not applicable

## Availability of data and materials

Sequence files are available at NCBI SRA under accession number SRR8468286-9.

## Authors’ contributions

BGDR and VW conceptualized and supervised the whole study. BGDR and XS interpreted the data and co-wrote the manuscript. MS, DB and VW performed all the microbial works and chemical treatments and prepared all the samples for RNA-Seq libraries. AK designed the RNA-Seq experiments and assembled the Illumina sequence reads. XS, MS and NBRK performed the biological interrogation and analysis of the RNA-Seq data. XS and NBRK performed all bio-computing works, statistical analyses, and genetic network modeling.

## Acknowledgments

We thank the ROIS supercomputing facilities of the National Institute of Genetics (Mishima, Japan) for the access to their RNA-Seq assembly pipeline for the assembly of the bacterial transcriptomic data generated in this study.

## Additional files

**Additional file 1: Figure S1** Scatter plots of the RPKM values for each transcriptome library of *E. coli* **(A)** and *L. monocytogenes* **(B)** in response to 1, 2 and 3 μg of O_3_ per gram of ripen fruits at 1 hour (hr), 2 hr and 3 hr, respectively, including the control and pure culture. (PowerPoint 521 KB)

**Additional file 2: Table S1 Sheet 1** Significance (up or down-regulated based on Pearson Correlation Coefficient), propensity and RPKM of *E. coli* genes during treatment of 1, 2, and 3 μg of O3 after 1, 2, and 3 hour (hr). **Sheet 2** Significance (up or down-regulated based on Pearson Correlation Coefficient), propensity and RPKM of *L. monocytogenes* genes during treatment of 1, 2, and 3 μg of O3 after 1, 2, and 3 hour (hr). The clusters in columns AN and AQ were shown in Fig. 4. (XLSX 1,139 KB)

**Additional file 3: Table S2 Sheet 1** Propensity of *E. coli* genes selected for 1 μg O_3_ network analysis. The 250 most highly expressed and 250 most lowly expressed genes from each treatment [control, 1, 2, and 3 hour (hr)]. Label numbers from 0 to 19 denote genes from most lowly expressed to most highly expressed. Only those fall into either label 0 or 19 at least in one treatment were selected. Genes differentially expressed based on Pearson Correlation Coefficient (PCC) were shown in columns L-N. **Sheet 2** Propensity of *E. coli* genes selected for 2 μg O_3_ network analysis. The 250 most highly expressed and 250 most lowly expressed genes from each treatment [control, 1, 2 and 3 hour (hr)]. Label numbers from 0 to 19 denote genes from most lowly expressed to most highly expressed. Only those fall into either label 0 or 19 at least in one treatment were selected. Genes differentially expressed based on PCC were shown in columns L-N. **Sheet 3** Propensity of *E. coli* genes selected for 3 μg O_3_ network analysis. The 250 most highly expressed and 250 most lowly expressed genes from each treatment [control, 1, 2 and 3 hour (hr)]. Label numbers from 0 to 19 denote genes from most lowly expressed to most highly expressed. Only those fall into either label 0 or 19 at least in one treatment were selected. Genes differentially expressed based on Pearson Correlation Coefficient were shown in columns L-N. **Sheet 4** Co-expression network of *E. coli* genes for 1 μg O_3_ at 1 hr. ‘color’: ‘brown’ demotes positively correlated by PCC; ‘color’: ‘green’ demotes negatively correlated by PCC. **Sheet 5** Co-expression network of *E. coli* genes for 1 μg O_3_ at 2 hr. ‘color’: ‘brown’ demotes positively correlated by PCC; ‘color’: ‘green’ demotes negatively correlated by PCC. **Sheet 6** Co-expression network of *E. coli* genes for 1 μg O_3_ at 3 hr. ‘color’: ‘brown’ demotes positively correlated by PCC; ‘color’: ‘green’ demotes negatively correlated by PCC. **Sheet 7** Co-expression network of *E. coli* genes for 1 μg O_3_ control. ‘color’: ‘brown’ demotes positively correlated by PCC; ‘color’: ‘green’ demotes negatively correlated by PCC. **Sheet 8** Co-expression network of *E. coli* genes for 2 μg O_3_ at 1 hr. ‘color’: ‘brown’ demotes positively correlated by PCC; ‘color’: ‘green’ demotes negatively correlated by PCC. **Sheet 9** Co-expression network of *E. coli* genes for 2 μg O_3_ at 2 hr. ‘color’: ‘brown’ demotes positively correlated by PCC; ‘color’: ‘green’ demotes negatively correlated by PCC. **Sheet 10** Co-expression network of *E. coli* genes for 2 μg O_3_ at 3 hr. ‘color’: ‘brown’ demotes positively correlated by PCC; ‘color’: ‘green’ demotes negatively correlated by PCC. **Sheet 11** Co-expression network of *E. coli* genes for 2 μg O_3_ control. ‘color’: ‘brown’ demotes positively correlated by PCC; ‘color’: ‘green’ demotes negatively correlated by PCC. **Sheet 12** Co-expression network of *E. coli* genes for 3 μg O_3_ at 1 hr. ‘color’: ‘brown’ demotes positively correlated by PCC; ‘color’: ‘green’ demotes negatively correlated by PCC. **Sheet 13** Co-expression network of *E. coli* genes for 3 μg O_3_ at 2 hr. ‘color’: ‘brown’ demotes positively correlated by PCC; ‘color’: ‘green’ demotes negatively correlated by PCC. **Sheet 14** Co-expression network of *E. coli* genes for 3 μg O_3_ at 3 hr. ‘color’: ‘brown’ demotes positively correlated by PCC; ‘color’: ‘green’ demotes negatively correlated by PCC. **Sheet 15** Co-expression network of *E. coli* genes for 3 μg O_3_ control. ‘color’: ‘brown’ demotes positively correlated by PCC; ‘color’: ‘green’ demotes negatively correlated by PCC. (XLSX 1,055 KB)

**Additional file 4: Table S3 Sheet 1** Propensity of *L. monocytogenes* genes selected for 1 μg O_3_ network analysis. The 250 most highly expressed and 250 most lowly expressed genes from each treatment [control, 1, 2, and 3 hour (hr)]. Label numbers from 0 to 19 denote genes from most lowly expressed to most highly-expressed. Only those fall into either label 0 or 19 at least in one treatment were selected. Genes differentially expressed based on Pearson Correlation Coefficient (PCC) were shown in columns L-N. **Sheet 2** Propensity of *L. monocytogenes* genes selected for 2 μg O_3_ network analysis. The 250 most highly expressed and 250 most lowly expressed genes from each treatment [control, 1, 2 and 3 hour (hr)]. Label numbers from 0 to 19 denote genes from most lowly expressed to most highly expressed. Only those fall into either label 0 or 19 at least in one treatment were selected. Genes differentially expressed based on PCC were shown in columns L-N. **Sheet 3** Propensity of *L. monocytogenes* genes selected for 3 μg O_3_ network analysis. The 250 most highly expressed and 250 most lowly expressed genes from each treatment [control, 1, 2 and 3 hour (hr)]. Label numbers from 0 to 19 denote genes from most lowly expressed to most highly expressed. Only those fall into either label 0 or 19 at least in one treatment were selected. Genes differentially expressed based on Pearson Correlation Coefficient were shown in columns L-N. **Sheet 4** Co-expression network of *L. monocytogenes* genes for 1 μg O_3_ at 1 hr. ‘color’: ‘brown’ demotes positively correlated by PCC; ‘color’: ‘green’ demotes negatively correlated by PCC. **Sheet 5** Co-expression network of *L. monocytogenes* genes for 1 μg O_3_ at 2 hr. ‘color’: ‘brown’ demotes positively correlated by PCC; ‘color’: ‘green’ demotes negatively correlated by PCC. **Sheet 6** Co-expression network of *L. monocytogenes* genes for 1 μg O_3_ at 3 hr. ‘color’: ‘brown’ demotes positively correlated by PCC; ‘color’: ‘green’ demotes negatively correlated by PCC. **Sheet 7** Co-expression network of *L. monocytogenes* genes for 1 μg O_3_ control. ‘color’: ‘brown’ demotes positively correlated by PCC; ‘color’: ‘green’ demotes negatively correlated by PCC. **Sheet 8** Co-expression network of *L. monocytogenes* genes for 2 μg O_3_ at 1 hr. ‘color’: ‘brown’ demotes positively correlated by PCC; ‘color’: ‘green’ demotes negatively correlated by PCC. **Sheet 9** Co-expression network of *L. monocytogenes* genes for 2 μg O_3_ at 2 hr. ‘color’: ‘brown’ demotes positively correlated by PCC; ‘color’: ‘green’ demotes negatively correlated by PCC. **Sheet 10** Co-expression network of *L. monocytogenes* genes for 2 μg O_3_ at 3 hr. ‘color’: ‘brown’ demotes positively correlated by PCC; ‘color’: ‘green’ demotes negatively correlated by PCC. **Sheet 11** Co-expression network of *L. monocytogenes* genes for 2 μg O_3_ control. ‘color’: ‘brown’ demotes positively correlated by PCC; ‘color’: ‘green’ demotes negatively correlated by PCC. **Sheet 12** Co-expression network of *L. monocytogenes* genes for 3 μg O_3_ at 1 hr. ‘color’: ‘brown’ demotes positively correlated by PCC; ‘color’: ‘green’ demotes negatively correlated by PCC. **Sheet 13** Co-expression network of *L. monocytogenes* genes for 3 μg O_3_ at 2 hr. ‘color’: ‘brown’ demotes positively correlated by PCC; ‘color’: ‘green’ demotes negatively correlated by PCC. **Sheet 14** Co-expression network of *L. monocytogenes* genes for 3 μg O_3_ at 3 hr. ‘color’: ‘brown’ demotes positively correlated by PCC; ‘color’: ‘green’ demotes negatively correlated by PCC. **Sheet 15** Co-expression network of *L. monocytogenes* genes for 3 μg O_3_ control. ‘color’: ‘brown’ demotes positively correlated by PCC; ‘color’: ‘green’ demotes negatively correlated by PCC. (XLSX 4,992 KB)

